# Boosting serotonin increases information gathering by reducing subjective cognitive costs

**DOI:** 10.1101/2021.12.08.471843

**Authors:** Jochen Michely, Ingrid M. Martin, Raymond J. Dolan, Tobias U. Hauser

## Abstract

Serotonin is implicated in the valuation of aversive costs, such as delay or physical effort. However, its role in governing sensitivity to cognitive effort, for example deliberation costs during information gathering, is unclear. We show that week-long treatment with a serotonergic antidepressant enhances a willingness to gather information when trying to maximize reward. Using computational modelling, we show this arises from a diminished sensitivity to subjective deliberation costs during the sampling process. This result is consistent with the notion that serotonin alleviates sensitivity to aversive costs in a domain-general fashion, with implications for its potential contribution to a positive impact on motivational deficits in psychiatric disorders.

## Introduction

Harvesting information about the world is essential for successfully navigating it. However, gathering information is costly and we need to balance between gathering too little and too much information (Gottlieb *et al*., 2013). Excess information gathering for a simple or irrelevant decision, such as seeking a homeopathic remedy for COVID-19, leads to unnecessary costs, including the time and energy spent exploring. Alternatively, expending too few resources on information gathering about an important decision, such as whether to undergo surgery, may result in considerable long-term aversive outcomes if the decision turns out to be ill-advised.

This challenge in information gathering can be reformulated as an arbitration between the value of novel information and the cognitive costs incurred. Humans struggle with solving this arbitration optimally, often showing excessive or insufficient information gathering (Bogacz *et al*., 2010; Hauser *et al*., 2017b). Interestingly, suboptimalities in information gathering are a feature in a range of psychiatric disorders (Clark *et al*., 2006; Taylor Tavares *et al*., 2007; Moutoussis *et al*., 2011; Hauser *et al*., 2017a; Hauser *et al*., 2017b; Ermakova *et al*., 2019).

The neurocomputational mechanisms underlying this arbitration between a putative gain, such as reward, and the associated costs, such as allocation of cognitive resources, remain unclear. It is hypothesised that the neurotransmitter serotonin may play a role in this process (Husain & Roiser, 2018). Even though the exact impact of serotonin on decision-making remains somewhat elusive (Cools *et al*., 2011; Dayan, 2012), it has been suggested as signalling a cost related to action, such as physical effort (Meyniel *et al*., 2016) or action inhibition (Crockett *et al*., 2009; Guitart-Masip *et al*., 2014).

Meyniel *et al*., 2016 demonstrated that boosting serotonin by selective serotonin reuptake inhibitors (SSRIs) reduces a perception of physical effort by lowering a sensitivity to its associated aversive costs. Likewise, it has been hypothesised that serotonin’s impact on intertemporal choice may reflect a reduced cost perception for delay-induced costs (Schweighofer *et al*., 2008; Miyazaki *et al*., 2014; Fonseca *et al*., 2015). However, it remains uncertain whether serotonin signals a cost beyond mere physical effort, for example the cost of cognitive effort critical for deliberation and information gathering (Froböse & Cools, 2018; Petitet *et al*., 2020).

In this study, we tested whether serotonin alleviates cognitive cost sensitivity in information gathering. Using a double-blind, placebo-controlled, between-subjects design, we assessed the impact of week-long daily treatment with the SSRI citalopram on decision-making during an established sequential information gathering task (Hauser *et al*., 2017a; Hauser *et al*., 2017b). We find SSRI treatment boosts information gathering in a manner indicative of subjects being more willing to exert cognitive effort in order to obtain reward. Computational modelling revealed this serotonergic effect is specifically driven by a reduction in the aversive cost of deliberation. The findings are consistent with serotonin playing a domain-general role in encoding ongoing both physical and cognitive costs, a mechanism that might be relevant for therapeutic approaches to motivational deficits in psychiatric disorders.

## Results

### Number of draws as a reliable measure of information gathering

In a double-blind, placebo-controlled, between-subjects pharmacological study, we tested 66 healthy volunteers who received a daily oral dose of the SSRI citalopram (20mg) or placebo over a period of seven consecutive days. To examine effects of acute and prolonged SSRI administration, we tested information gathering on two occasions, once after the first dose on day 1, and once after week-long treatment on day 7.

In both sessions, subjects performed a modified version of an information sampling task (Clark *et al*., 2006; Hauser *et al*., 2017a; Hauser *et al*., 2017b). On each game, subjects saw 25 covered cards (Fig. 1A, grey squares) and had to decide which of the two distinct colours formed the overall majority (colours varied across games). Initially, all cards were covered, and subjects were allowed to open as many cards as they wished before committing to a decision. To alter the incentive for information gathering, subjects were exposed to two distinct reward conditions. In the ‘fixed’ condition, gathering information did not come incur a monetary cost (subjects received 100 points for correct decisions, independent of the number of sampled cards prior to the decision). In the ‘decreasing’ condition, gathering information came with an explicit monetary cost, where opening each card led to a 10-points reduction in potential wins. Subjects started from a maximum potential gain of 250 points so that, for example, a correct decision after seven draws would lead to a gain of 180 points (250-7*10=180). In both conditions, incorrect decisions resulted in a loss of 100 points, independent of the amount of prior sampling.

**Figure 1.**
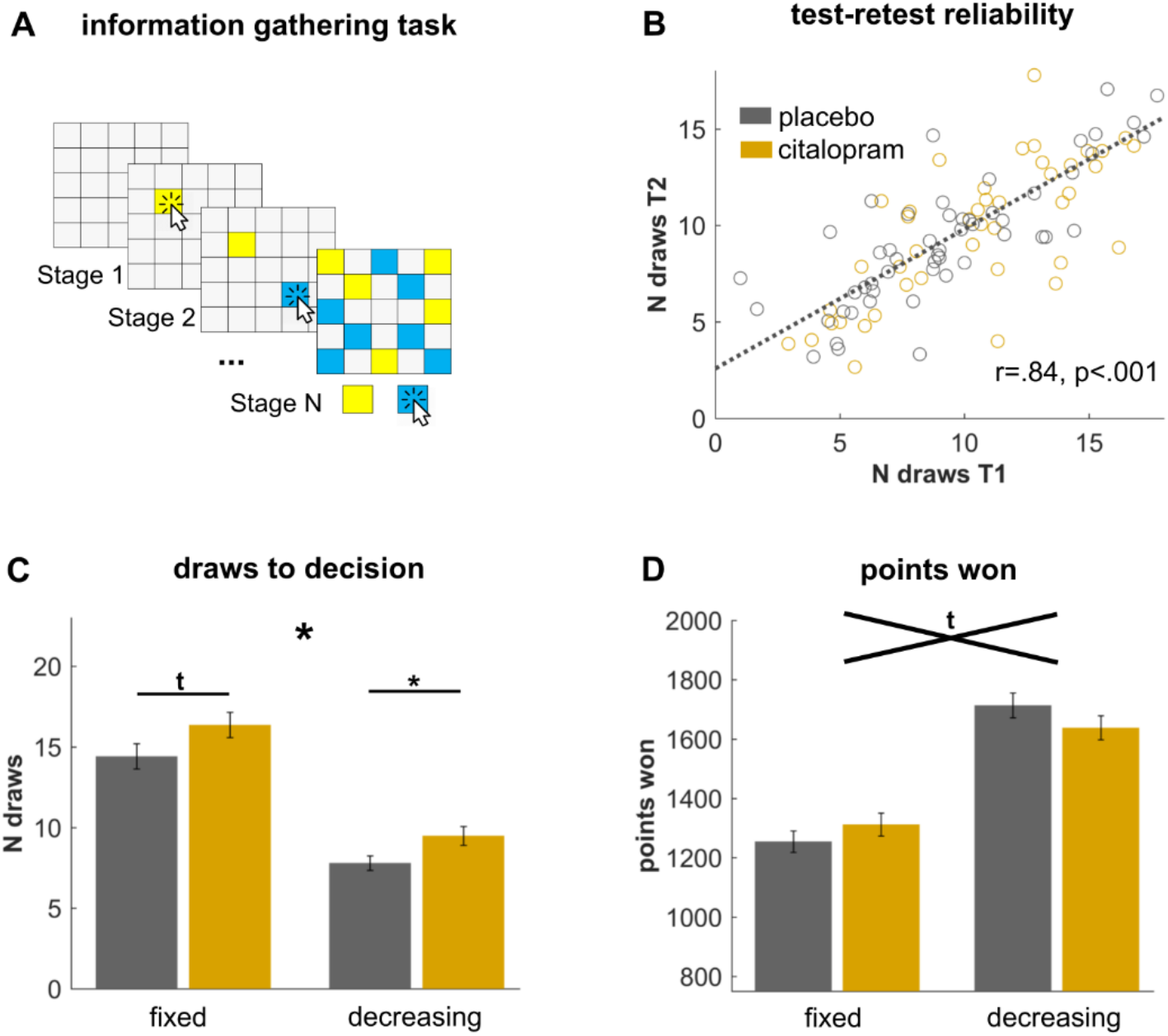
Increased information gathering after SSRI administration. (A) Subjects were randomly allocated to a daily dose of 20mg citalopram or placebo for seven consecutive days. Subjects performed the information gathering task on two sessions: session 1 took place on day 1 after a single-dose administration, session 2 took place on day 7 after week-long drug treatment. For each game, subjects started with a fully covered deck of 25 grey cards. They were allowed to open as many cards as they wished before declaring a decision about whether they believed the majority of cards was yellow or blue (colours varied across games). In the ‘fixed’ condition, no external costs for sampling applied, but in a ‘decreasing’ condition a potential total win of 250 points reduced by 10 points per uncovered card. In both conditions, subjects lost 100 points upon making an incorrect decision. (B) Information gathering (number of draws) shows a high test-retest reliability, demonstrating that the task is capable of reliably detecting individual differences in information gathering. (C) SSRIs led to a general increase in the number of draws needed before declaring a decision (shown here collapsed across both timepoints as there were no interactions with time). This was evident across both conditions, but was particularly pronounced in the decreasing condition. (D) The SSRI-induced increase in sampling resulted in SSRI subjects winning (marginally) more points in the fixed (where sampling is cost-free), and less points in the decreasing (where sampling is costly) condition, as compared to placebo subjects. * p < 0.05, ^t^ p < 0.1. Error bars indicate SEM.

We assessed the number of draws a subject made before committing to a decision as a key indicator of information gathering in the task. This metric is a proxy for a subject’s need to gather information, and has been found sensitive to individual differences including levels of psychopathology (Clark *et al*., 2006; Taylor Tavares *et al*., 2007; Moutoussis *et al*., 2011; Hauser *et al*., 2017a; Hauser *et al*., 2017b).

A potential limitation with this metric – as with many other cognitive variables (Shahar *et al*., 2019) – is that little is known about its psychometric properties, such as test-retest reliability. Good psychometric properties are particularly important when tasks are used to make inference across repeated measurements, or when used to differentiate between subjects (e.g. patient vs non-patient groups, drug vs no-drug groups). Given that we assessed the same subjects twice, with a 7-day interval between measurements, we were in the unique position to assess the test-retest reliability of the main measure of interest, the number of draws. We found that the number of draws was remarkably stable across sessions (both groups: r=0.84, p<0.001; placebo group alone: r=0.88, p<0.001; Fig. 1B), indicative of high reliability at par with well-established and validated psychometric questionnaire measures (Fleiss, 1986; Cicchetti, 1994). This means that our information gathering task is well-suited for studying individual differences.

### Serotonin increases information gathering

To assess the effects of our pharmacological intervention, we next compared the number of draws between drug groups using a repeated-measures ANOVA with within-subject factors drug (placebo, SSRI) and time (single-dose, week-long). We found a significant main effect of drug (F_1,64_=5.0, p=0.028; Fig. 1C), driven by an increase in number of draws in the SSRI compared to placebo subjects. Further analyses showed that this was particularly prominent in the decreasing condition when sampling information was costly (decreasing: t_64_=2.3, p=0.025; fixed: t_64_=1.8, p=0.084).

Additionally, we found a significant effect of condition (F_1,64_=202.3, p<0.001) but no interaction with drug (F_1,64_=0.07, p=0.793), indicating that subjects gathered more information in the fixed compared to the decreasing condition across the entire sample. The duration of drug administration (single-dose vs. week-long) did not impact the drug’s effect on information gathering (drug x time: F_1,64_=0.9, p=0.345), or the interaction with condition (drug x time x condition: F_1,64_=0.3, p=0.602). This means that the drug effects were similar for acute and prolonged treatment. Overall, these results show that (acute and week-long) serotonergic treatment enhanced a willingness to gather more information before declaring a choice.

### Serotonergic effects on further task metrics

Next, we assessed whether SSRIs affected other metrics, not directly related to information gathering. First, we examined subjects’ accuracy, as measured by whether a subject chooses the colour that is more plentiful at the time of decision. We did not find any pharmacological effect on accuracy, suggesting that SSRIs do not simply change motivation, attention or information processing in general (drug: F_1,64_=0.1, p=0.711; drug x time: F_1,64_=0.7, p=0.402; drug x condition: F_1,64_=0.4, p=0.537; drug x time x condition: F_1,64_=0.5, p=0.482; Supplementary Fig. S1).

Lastly, we investigated pharmacological effects on task earnings, i.e. how many points subjects won in the task. This measure is a conglomerate measure influenced by information gathering, accuracy, and luck. When assessing how drug treatment affected task gains, we found a statistical trend for a drug by condition interaction (F_1,64_=3.6, p=0.060; Fig. 1D). This reflects an SSRI-induced increase in sampling across both conditions, as enhanced information gathering in the fixed condition (where sampling is cost-free) leads to an increase in earnings, whereas an oversampling in the decreasing condition (where sampling is costly) leads to a decrease in earnings (Hauser *et al*., 2017b). This result suggests that SSRI treatment increased information gathering per se, independent of condition, and thus independent of reward maximisation.

### Emerging subjective costs reduce information gathering

To decipher the computational mechanisms that drive an increase in information gathering in SSRI-treated subjects, we fitted three different computational models to individual subjects’ choices (cf. Methods). These previously developed and evaluated models cast information gathering as a trade-off between information gain of a new sample and the incurred costs in sampling information. In particular, they capture subjective information gathering costs in the context of Bayesian decision making and characterise how these costs change as information gathering continues (Hauser *et al*., 2017a; Hauser *et al*., 2017b; Bowler *et al*., 2021).

In line with our previous findings (Hauser *et al*., 2017a; Hauser *et al*., 2017b; Bowler *et al*., 2021), the winning model revealed that the costs for gathering information is subjective and changes over the course of information gathering. Model comparison revealed that these costs were not represented as per explicit instruction (‘objective’ model; i.e. no costs for the fixed, and -10 for the decreasing condition), but that ‘subjective’ costs accumulated as information gathering continued, meaning that it becomes subjectively more and more costly to gather further information as a function of cumulative information gathering. This effect was best captured by a model in which costs escalate in a ‘non-linear’, rather than a ‘linear’, fashion (Fig. 2A), in accordance with previous studies which identified the same winning model (Hauser *et al*., 2017a; Hauser *et al*., 2017b; Hauser *et al*., 2018; Bowler *et al*., 2021).

**Figure 2.**
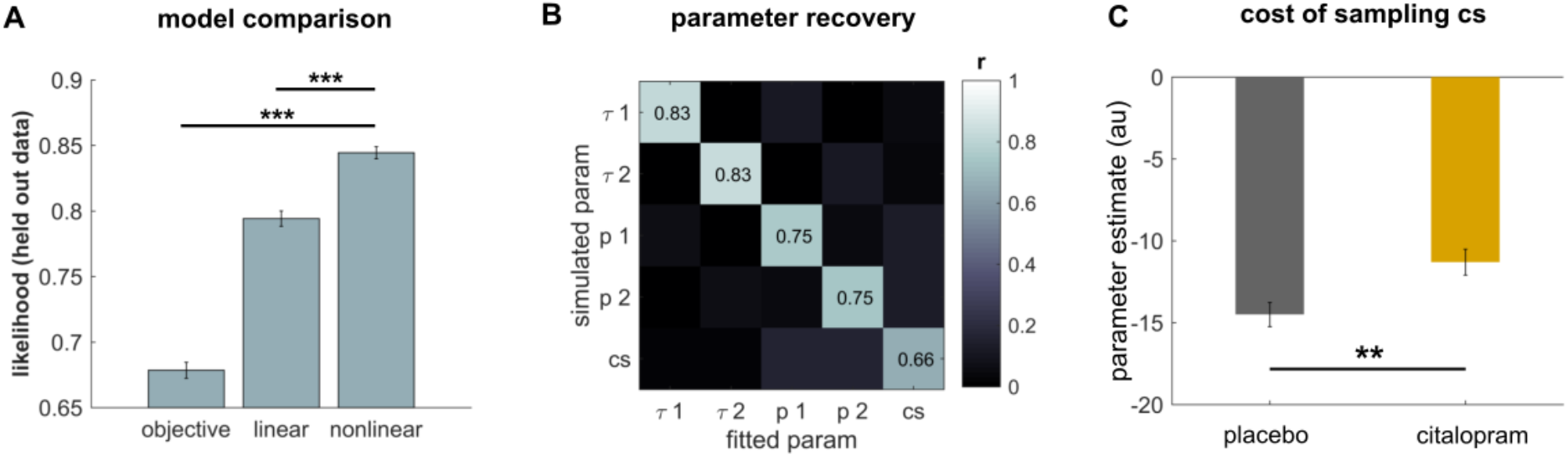
Computational modelling. (A) Model comparison predicting cross-validated hold-out data revealed that a nonlinear increase in subjective sampling costs fitted subjects’ performance best. (B) Confusion matrices of the winning model shows that the parameters could be recovered using 50’000 simulated agents. This was demonstrated by medium to large correlations of the fitted with the original parameters used for simulation. 1: fixed condition, 2: decreasing condition. (C) SSRI reduce subjective costs in information gathering, indicated by a lower subjective cost model parameter in citalopram as compared to placebo subjects (collapsed across both time points as there was no time effect on result). *** p < 0.001, ** p < 0.01, * p < 0.05. Error bars indicate SEM.

When we investigated how serotonin impacted putative mechanisms underlying information gathering, using simulations, we found that model parameter estimates could be accurately recovered, demonstrated in a positive association between parameter estimates derived from fitting to real and simulated data (Fig. 2B).

### SSRI decreases subjective costs of information gathering

To better understand how serotonin impacted information gathering, we compared the model parameters between the two groups. We found that only one model parameter that survived multiple comparison correction (cf. Supplementary Fig. S2). In particular, we found that SSRI treatment specifically affected the subjective cost parameter *cs* (F_1,64_=7.8, p=0.007, uncorr.; p=0.042, Bonferroni-corr.; Fig. 2C). This parameter is less negative in the SSRI group, which means that SSRI treatment led to a reduction in the subjective costs of information gathering. In alignment with the behavioural findings, there was no effect of time (acute vs week-long treatment) on this parameter (drug x time interaction: F_1,64_=.056, p=.814), meaning that acute and week-long treatment had a similar effect. In sum, this indicates that an increase in information gathering in SSRI-treated subjects arises as a consequence of a lowered sensitivity to the subjective cost of sampling after serotonergic intervention.

## Discussion

We show that boosting serotonin function pharmacologically leads to increased information gathering in a sequential information sampling task. Using computational modelling, we demonstrate this effect arises from a serotonin-induced reduction in subjective sampling costs.

Serotonin is an impactful neuromodulator, but its precise influence on cognition, motivation and behaviour remains elusive (Dayan, 2012; Olivier, 2015). Whilst early accounts proposed serotonin as an opponent to dopamine that is mainly involved in signalling aversive outcomes (Daw *et al*., 2002; Cools *et al*., 2011), recent evidence has drawn a more complex and nuanced picture. In particular, studies suggest serotonin plays a crucial role in signalling the costs that are associated with aversive experience. These costs encompass punishment or delay (Denk *et al*., 2005; Cools *et al*., 2008; Schweighofer *et al*., 2008; Tanaka *et al*., 2009; den Ouden *et al*., 2013; Miyazaki *et al*., 2014), and extend to costs associated with the exertion of physical effort (Meyniel *et al*., 2016). Our finding confirms and expands this theory by demonstrating that enhancing central serotonin also modifies a subjective perception of cognitive effort costs in the context of information gathering.

These subjective costs sit right at the heart of information gathering. Over recent years, a growing literature spanning non-human animal neurophysiology and computational modelling of human behaviour demonstrates the relevance of subjective costs in the context of sequential sampling tasks (Thura & Cisek, 2016; Hauser *et al*., 2017b). In essence, mounting subjective costs means that subjects become less willing to sample more information as they go along, and are drawn towards making a decision in the absence of clear evidence. Neurophysiologically, it has been suggested this process may be implemented by a neural signal that increases over time, often termed urgency signal (Cisek *et al*., 2009; Yau *et al*., 2020). This urgency signal is added to an accumulation of evidence in (pre-)motor areas, thus promoting decisions before absolute evidence is being gathered (Thura & Cisek, 2016).

In our computational model, we capture this behavioural signature by means of subjective sampling costs that are imposed on the action value for non-deciding, thus making a continuation of gathering novel information less and less likely. Replicating previous findings, we show such cognitive costs are low in the beginning but escalate in a non-linear manner as sampling continues over time. This non-linear emergence of costs can help explain suboptimalities in human behaviour, with an undersampling in cost-free and an oversampling in costly environments (Bogacz *et al*., 2010; Hauser *et al*., 2017b).

In this study, we show that serotonin specifically reduces the scaling of such costs, meaning that subjective costs are impacting a decision less, irrespective of how much one has already sampled. This finding has putative clinical relevance. SSRIs constitute a first-line intervention in the treatment of depression (Cipriani *et al*., 2018), a disorder typically characterised by motivational deficits, such as lowered willingness to engage in effortful behaviour (Der-Avakian *et al*., 2016). Critically, using similar laboratory tasks, studies have shown that a willingness to gather information is reduced in patients suffering from depression (Taylor Tavares *et al*., 2007), as well as healthy individuals with low levels of motivation (Roets *et al*., 2008). Our study suggests that SSRI treatment can counteract such motivational deficits, through an increase in information gathering that is mediated by reduced cost perception. This result nicely extends previous computational studies that revealed how SSRI intervention increases physical effort exertion via reduced perception of effort costs (Meyniel *et al*., 2016). Taken together, these findings suggest that serotonin is not only relevant in signalling the costs of physical, but also of mental effort, hinting at a domain-general role of serotonin in overcoming aversive costs. Overall, this mechanism may help explain how serotonergic treatment gives rise to an alleviation of motivational deficits in patients suffering from depression.

More speculative is a link between our current finding and another psychiatric disorder, obsessive-compulsive disorder (OCD), that has been linked to aberrant information processing (Volans, 1976; Fear & Healy, 1997; Pelissier & O’Connor, 2002). Studies show (Hauser *et al*., 2017b; Voon *et al*., 2017), although not consistently (Chamberlain *et al*., 2007; Jacobsen *et al*., 2012), excessive information gathering behaviour in OCD patients. Our findings suggest that SSRIs, first-line agents for treatment of OCD, could further exacerbate information gathering, although previous studies did not find an association with medication status (Hauser *et al*., 2017b). It is possible, however, that baseline serotonin levels may play a critical role in modulating an impact of serotonergic medication on information gathering. For instance, the relationship between serotonin levels and information gathering may follow an inverted u-shape function, which could also explain why a reduction of serotonin levels by means of tryptophan depletion in a previous study induced a change in sampling in a similar direction as the SSRI effects revealed in the current study (Crockett *et al*., 2012).

Lastly, we found that serotonin increased information gathering per se, independent of the experimental condition, thereby increasing earnings of SSRI subjects in a cost-free, and reducing earnings in a costly environment, where sampling comes at a small cost. Note that this suggests information gathering was enhanced independent of reward maximisation, as SSRI subjects were willing to pay small, local costs for more information. On the other hand, however, this indicates subjects were more averse to experiencing large, global costs, as, in both conditions, a greater amount of information makes a wrong decision, and thereby a large monetary loss, less likely. Critically, this effect is in line with influential theories of serotonin function postulating its prominent role in the avoidance of aversive outcomes (Dayan & Huys, 2008).

In conclusion, our findings demonstrate that SSRI administration enhances information gathering. Computational modelling indicated this arises from a reduced perception of subjective sampling costs, rendering information gathering less cognitively effortful. Our findings point to a candidate mechanism by which serotonergic treatment might help alleviate motivational deficits in the context of a range of mental illnesses.

## Methods

### Subjects

66 healthy volunteers (mean age: 24.7±3.9; range 20-38 years; 40 females; Supplementary Table S1) participated in this double-blind, placebo-controlled study. All subjects underwent an electrocardiogram to exclude QT interval prolongation and a thorough medical screening interview to exclude any neurological or psychiatric disorder, any other medical condition, or medication intake. The experimental protocol was approved by the University College London (UCL) local research ethics committee, with informed consent obtained from all participants. Data from different tasks of the same participants were already published elsewhere (Michely *et al*., 2020a; Michely *et al*., 2020b).

### Pharmacological procedure

Participants were randomly allocated to receive a daily oral dose of the SSRI citalopram (20mg) or placebo, over a period of seven consecutive days. All subjects performed two laboratory testing sessions. The first session was on day 1 of treatment, approximately 3h after single dose administration, as citalopram reaches its highest plasma levels after this interval (Noble & Benfield, 1997). On the following days, subjects were asked to take their daily medication dose at a similar time of day, either at home or at the study location. The second session was on day 7 of treatment, with the tablet being taken approximately 3h before the experiment.

### Experimental task

We examined sequential information gathering using a modified version of an information sampling task (Clark *et al*., 2006; Hauser *et al*., 2017a; Hauser *et al*., 2017b). On each game, subjects saw 25 covered cards (Fig. 1A, grey squares) and had to decide whether the majority of cards was of colour 1 or colour 2 (e.g., yellow or blue, colours varied across games). Using a computer mouse, subjects were allowed to sample as many cards as they wished before committing to one of the two colours.

The first 15 games were part of a “fixed” condition, in which gathering additional information was not costly. Specifically, subjects received 100 points for correct decisions and lost 100 points for incorrect decisions, irrespective of the number of cards opened or the time spent on task before decision. In the “decreasing” condition, information gathering incurred external costs resulting in a reduction of potential gains. Specifically, starting from a maximum potential gain of 250 points, opening each card led to a 10-point gain reduction (e.g., gain after 7 opened cards: 250 - 7 * 10 = 180 points). Incorrect decisions resulted in a loss of 100 points, independent of the amount of prior sampling.

After each game, subjects were informed about their gains, and then proceeded to the next game. The game sequences were selected so that 10 games in each condition were relatively difficult with a generative probability close to 50% (similar to that in the original information sampling task; (Clark *et al*., 2006). An additional 5 sequences were easier with a clearer majority (generative probabilities of a binomial process p ∼ 0.7) to allow for a broader variability in information gathering. Order of sequences was randomized. Before the first game, subjects performed a practice game to familiarize themselves with the task.

### Computational modelling

To investigate the computational mechanisms underlying information gathering, we used a computational model that we have previously developed for this task. Here, we reiterate the most relevant equations, but a detailed description of the model can be found in (Hauser *et al*., 2017b) and subsequent papers (Hauser *et al*., 2017a; Hauser *et al*., 2018; Bowler *et al*., 2021).

The computational model assumes that agents make inference about which colour is more likely to form the majority of cards *P*(*MY* | *n*_*y*_, *N*) with *MY* being a majority of yellow, given the current amount of yellow cards (*n*_*y*_) out of a total of *N* sampled cards. *P(MY)* is calculated using the current number of cards, making inference about the generative probability that could have caused this distribution of cards (cf. (Hauser et al., 2017b)).

This belief is then used calculate the action *Q* values for declaring for yellow and blue, weighting both potential gains and losses by the inferred likelihood of them taking place

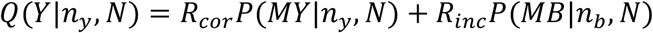

with *R*_*cor*_ and *R*_*inc*_ being the potential wins and losses (set to +100 / -100 here; for discussion cf. (Hauser et al., 2018)).

A more challenging computation is the estimation of the action value for not deciding (*Q(ND)*). This is computed as the sum of the value of the future states (*V(s’)*, a weighted sum of the *Q* values in that state), weighted by how likely they will materialise (*p(s’*|*n*_*y*_,*N)*; i.e. how likely will I end up in that state given the cards that I have opened so far). In addition, a cost per step *c* incurs, which accounts for the subjective costs for sampling more information and thus spending more effort, time, and points (in the decreasing condition) on gathering information.

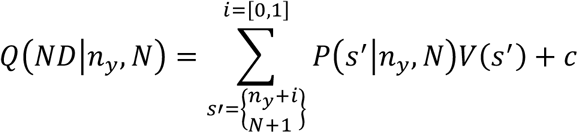

In accordance with previous studies, we found that subjective costs did not follow the explicit costs (i.e., 0 in the fixed, -10 points in the decreasing condition; ‘objective’ model). Moreover, costs also did not increase linearly (‘linear’ model), but rather scaled in a non-linear fashion during sampling (‘nonlinear’ model, modelled as a sigmoid). The best performing, non-linear, model comprised two free parameters, a scaling parameter *cs* that determined how big the maximal costs could be, and an intercept *p*, which determined after how many samples (*n*) these costs started to escalate (Drugowitsch *et al*., 2012; Murphy *et al*., 2016).

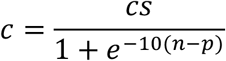

Lastly, the choice policy was determined using a softmax rule with decision temperature τ and an additional epsilon greedy element (ξ) that captures choices that were not adequately captured by the model. The policy was not only used for choice arbitration, but also in the planning process to inform state values und backward planning (cf. (Hauser *et al*., 2017b) for details):

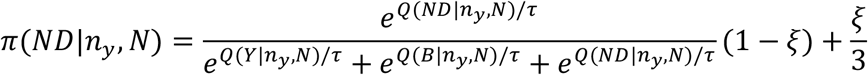

To determine the best-fitting model, we compared three distinct models with different forms of cost structures as described above: objective, linear, non-linear. To compare their model fit, we use out-of-sample prediction using 5-fold cross-validation assessing the predictive likelihood. This is a measure that finds an optimal balance between complexity and accuracy using the held-out data (Dubois *et al*., 2020). Model parameters were optimised using ‘fmincon’, with multiple starting points to overcome local minima.

### Statistical analysis

In this study, we tested whether SSRI treatment affects information gathering. The number of draws before a decision is a good indicator for the amount of information that a subject is willing to collect before committing to a decision (Hauser *et al*., 2017a; Hauser *et al*., 2017b). We analysed this behavioural metric using repeated-measures ANOVAs with the between-subject factor drug (SSRI, placebo), and the within-subject factor condition (fixed, decreasing). To assess whether effects were different after single-dose administration, compared with one-week treatment, we added the within-subject factor time (session 1: acute, session 2: week-long). Significant effects were further assessed using independent-sample t-tests (SSRI vs. placebo). As secondary measures (Hauser *et al*., 2017a; Hauser *et al*., 2017b), we assessed whether drug treatment affected how many points subjects won, and how accurate subjects were in their decision-making (i.e., how often subjects correctly opted for the colour with the current majority of cards), using the same statistical procedures. For comparison of computational model parameters, we applied Bonferroni correction for the number of model parameters.

### No drug effects on self-report questionnaires

To examine putative treatment effects on subjective affective states over the course of the study, participants completed the Beck’s Depression Inventory (BDI-II, (Beck *et al*., 1996)), Snaith-Hamilton Pleasure Scale (SHAPS, (Snaith *et al*., 1995)), State-Trait Anxiety Inventory (STAI, (Spielberger *et al*., 1983)), and the Positive and Negative Affective Scale (PANAS, (Watson *et al*., 1988)) on two different occasions: (i) pre-drug, day 1; (ii) peak drug, day 7. We found no evidence for serotonergic effects on any of the self-report questionnaires (Supplementary Table S1). This is in line with previous studies showing week-long SSRI treatment does not impact on mood in healthy volunteer participants (Harmer, 2013).

