## Supplementary Material for "Boosting serotonin increases information gathering by reducing subjective cognitive costs"

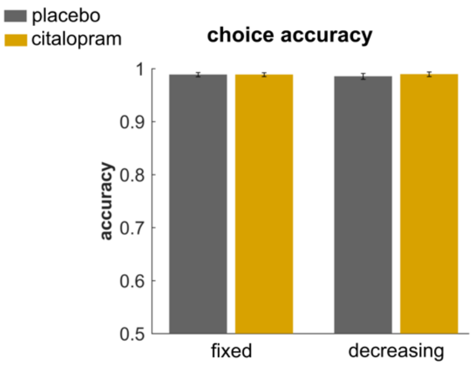

**Supplementary Figure S1**. We found no effect of drug or condition on choice accuracy (the probability of choosing the colour that currently forms the majority of cards at time of decision). This shows that SSRIs do not simply change motivation, attention or information processing in general. Error bars indicate SEM.

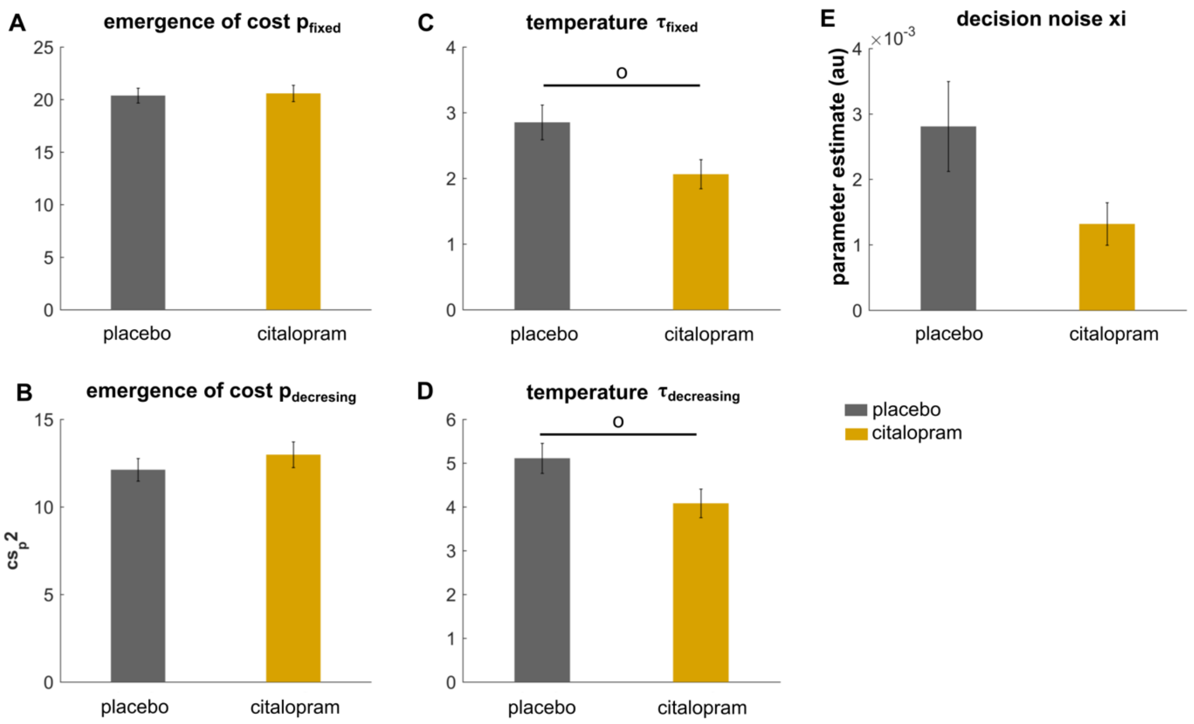

**Supplementary Figure S1**. We found that effect of drug on any of the other model parameters when correcting for multiple comparisons, using Bonferroni correction for six comparisons. ° $p < 0.1$ (uncorr.). Error bars indicate SEM.

| | placebo | | citalopram | | $P_{value}$ |
|---|---|---|---|---|---|
| gender | 20 ♀ / 13 ♂ | | 20 ♀ / 13 ♂ | | 1.000 |
| age | 24.8 ± 3.9 | | 24.5 ± 4.0 | | 0.757 |
| BDI – II [day 1] | 4.4 ± 5.4 | | 3.6 ± 4.0 | | 0.540 |
| BDI – II [day 7] | 4.6 ± 5.7 | | 4.5 ± 4.5 | | 0.924 |
| BDI – II [day 7 – day 1] | 0.2 ± 3.3 | | 0.8 ± 3.5 | | 0.469 |
| SHAPS [day 1] | 0.3 ± 1.0 | | 0.3 ± 0.7 | | 1.000 |
| SHAPS [day 7] | 0.6 ± 1.6 | | 0.8 ± 2.5 | | 0.771 |
| SHAPS [day 7 – day 1] | 0.3 ± 1.5 | | 0.5 ± 2.1 | | 0.738 |
| STAI - state [day 1] | 30.6 ± 8.5 | | 30.1 ± 6.4 | | 0.795 |
| STAI - state [day 7] | 33.1 ± 9.7 | | 31.4 ± 6.6 | | 0.392 |
| STAI - state [day 7 – day 1] | 2.5 ± 8.5 | | 1.4 ± 5.6 | | 0.508 |
| STAI - trait [day 1] | 33.1 ± 9.7 | | 34.6 ± 6.6 | | 0.479 |
| STAI - trait [day 7] | 34.6 ± 9.8 | | 35.5 ± 7.5 | | 0.664 |
| STAI - trait [day 7 – day 1] | 1.5 ± 5.0 | | 0.9 ± 3.1 | | 0.615 |
| PANAS - positive [day 1] | 31.2 ± 8.6 | | 30.0 ± 7.9 | | 0.562 |
| PANAS - positive [day 7] | 29.0 ± 10.4 | | 28.3 ± 8.3 | | 0.775 |
| PANAS - positive [day 7 – day 1] | -2.3 ± 7.3 | | -1.7 ± 5.6 | | 0.749 |
| PANAS - negative [day 1] | 11.5 ± 2.5 | | 11.2 ± 1.4 | | 0.588 |
| PANAS - negative [day 7] | 12.1 ± 3.3 | | 11.1 ± 1.7 | | 0.143 |
| PANAS - negative [day 7 – day 1] | 0.5 ± 3.2 | | -0.2 ± 1.8 | | 0.277 |

**Supplementary Table S2**. *Self-report questionnaire data.*
We found no impact of drug on any of self-report questionnaires. BDI – II = Beck's Depression Inventory II (Beck et al., 1996), SHAPS = Snaith-Hamilton Pleasure Scale (Snaith et al., 1995), STAI = State-Trait Anxiety Inventory (Spielberger, 1983), PANAS = Positive and Negative Affective Scale (Watson et al., 1988).
